# Mice immunized with N1 neuraminidase in a single cycle infectious virus or in a recombinant nanoparticle vaccine are protected against lethal challenge with highly pathogenic avian influenza H5N1 influenza

**DOI:** 10.64898/2026.10.11.758125

**Authors:** Michael S. Piepenbrink, Ramya S. Barre, Qinzhe Li, Ruby A. Escobedo, Ahmed M. Elsayed, Mark R. Walter, Jonathan F. Lovell, Luis Martinez-Sobrido, James J. Kobie

## Abstract

The recent spillover of highly pathogenic (HP) H5N1 clade 2.3.4.4b into dairy cattle and other mammalian species has been concerning. Vaccine candidates that can be rapidly synthesized and implemented are key countermeasures for protecting the human population against emerging pathogens. Here we examine the use of a pH1N1 virus lacking the genetic material for hemagglutinin (HA), limiting the infection to a single cycle (sciIAV) or Cobalt lipid-porphyrin nanoparticles decorated with neuraminidase (NA) of HPAI H5N1 (N1-CoPoP) as vaccine candidates. C57BL/6 mice that were inoculated intranasally with a single dose of pH1N1 sciIAV did not differ in serum NA antibody compared to uninoculated mice. However, mice that were inoculated intramuscularly with the N1-CoPoP alone or in combination with pH1N1 sciIAV had increased N1 IgG antibodies. Upon lethal challenge with HPAIV H5N1, all mice inoculated with either pH1N1 sciIAV or N1-CoPoP were protected from weight loss, unlike unvaccinated mice with substantial body weight losses that succumb to viral infection between 7 and 10 days post infection (dpi). Both pH1N1 sciIAV and N1-CoPoP immunization minimized viral infection such that only one mouse receiving pH1N1 sciIAV had detectable viral titer in the lungs at 4 dpi. By 6 dpi, no mouse inoculated with pH1N1 sciIAV and/or N1-CoPoP had detectable virus in nasal turbinates, lungs or the brain, unlike the high viral titers found in uninoculated mice. Both pH1N1 sciIAV and N1-CoPoP were protective and should be investigated further as alternative vaccine candidates.

## Introduction

Highly pathogenic avian influenza (HPAIV) H5N1 emerged from the A/goose/Guangdong/1/96 lineage first detected in domestic geese in China in 1996 and has since evolved into multiple clades, including the currently dominant clade 2.3.4.4b, which has driven an unprecedented global panzootic ^1^. HPAI H5N1 is now spread extensively by migratory wild birds and has reached every inhabited continent, causing substantial mortality in wild avian populations and repeated outbreaks in commercial poultry ^2,3^. Since 2021, the virus has expanded its host range dramatically, infecting numerous mammalian species including foxes, mink, bears, seals, sea lions, cats, and dairy cattle ^4^. In the United States, HPAI H5N1 was introduced through migratory birds and subsequently spread across wild bird populations, poultry operations, and, in the beginning of 2024, dairy cattle herds in multiple states, representing an unprecedented adaptation to a livestock mammalian host ^5^. Although sustained human-to-human transmission has not been observed, zoonotic infections have occurred following exposure to infected poultry and dairy cattle, highlighting the virus’s ability to cross species barriers ^6^. The continued circulation of HPAI H5N1 among diverse animal reservoirs increases opportunities for viral adaptation and reassortment, raising concerns about pandemic potential and underscoring the need for enhanced surveillance, preparedness, and broadly protective vaccines.

Single-cycle infectious influenza virus (sciIAV) represents a promising next-generation alternative to conventional inactivated influenza vaccines (IIV) because they combine the immunogenicity of a live viral infection with an improved safety profile ^7^. Unlike conventional vaccines, which primarily induce strain-specific antibody responses to surface antigens, sciIAV undergo only one round of infection because of mutations or deletions in a key component needed for infection or propagation and dissemination of new viruses. For instance, replacing the viral hemagglutinin (HA) gene with the sequence for the green fluorescent protein (GFP) results in viral progeny that lacks HA that can only be propagated in cells expressing the protein ^7–9^. In the absence of HA expressing cells, other viral proteins can be expressed but do not result in viral infectious progeny. This allows for stimulation of robust CD8⁺ T-cell responses, lung-resident memory T cells, neuraminidase (NA)-specific antibodies, and, in newer designs such as CLEARFLU, neutralizing HA antibodies, potentially providing both strain-matched protection and broader heterosubtypic immunity against diverse influenza viruses ^10^. These sciIAV vaccine platforms can elicit cross-reactive cellular immunity that is less susceptible to antigenic drift, a major limitation of current seasonal vaccines ^11,12^ while maintaining a favorable safety profile because viral replication is restricted to a single cycle. These characteristics make sciIAV vaccines attractive candidates for achieving broader and longer-lasting protection than conventional influenza vaccination approaches.

Current licensed influenza vaccines, primarily focus on inducing antibodies against the highly variable HA protein ^13^. NA is more antigenically conserved than HA ^14^, and NA-specific antibodies have been shown to independently correlate with protection against influenza infection and disease severity, providing broader immunity across drifted viral strains (reviewed in ^15,16^). Lipid nanoparticle (LNP)-based presentation of NA can further enhance immunogenicity by displaying the antigen in a highly ordered, multivalent format that promotes robust B cell activation and the generation of functional NA-inhibiting antibodies ^17^. Recent studies have demonstrated that NA-containing nanoparticle and liposome vaccine platforms induce stronger and broader humoral and cellular immune responses, improve protection against heterologous and heterosubtypic influenza viruses, and can complement conventional HA-based vaccines to expand cross-protection ^17,18^. Moreover, LNP technologies are compatible with modern mRNA and recombinant protein vaccine approaches, enabling rapid manufacturing and precise antigen design while overcoming the inconsistent NA content found in many current seasonal influenza vaccines ^19^. Collectively, NA-decorated LNP vaccines may provide more durable and broadly protective immunity than conventional influenza vaccines by targeting a more highly conserved viral antigen and enhancing its immunological presentation. Here we show that administering either sciIAV and/or N1 decorated cobalt lipid-porphyrin (N1-CoPoP) nanoparticles (N1-CoPoP) protect mice against HPAI H5N1 challenge.

## Results

Inoculating mice with pH1N1 sciIAV did not induce significant antibodies in serum toward N1 recombinant proteins, 40 days later (**Figures 1A, B, C**). This was expected, as sciIAV can infect cells but because the genetic material for HA is not present, progeny viruses cannot infect other cells. This, however, does not mean NA directed antibodies were not produced in sciIAV inoculated mice, but were at such a low concentration that levels were not different from that seen in mock treated mice. Intramuscular immunization of mice with CoPoP co-decorated with the N1 of A/Texas/37/2024 H5N1 and N1 A/Cambodia/NPH230032_2/2023 H5N1 (N1-CoPoP) on day 14 and 28 did induce antibodies in serum against N1 proteins by day 40 (**Figures 1A, B, C**). Serum IgG from these mice also had mild binding to N2 A/Darwin/09/2021 H3N2 (**Figure 1D**), but this could be accounted most likely by the use of the immunogenic tetramerization domain of human vasodilator-stimulated phosphoprotein (VASP) because binding to the monomer of N2 A/Darwin/09/2021 H3N2 was not significant (*p* > 0.05; **Figure 1E**) but binding to Nipah Virus glycoprotein containing the VASP tetramerization domain was significant (p < 0.0001) (**Figure 1F**) in serum from mice inoculated with N1-CoPoP alone or in combination with sciIAV. Only minimal IgG binding to H1 (Figure 1G) or H5 (**Figure 1H**) was observed.

**Figure 1.**
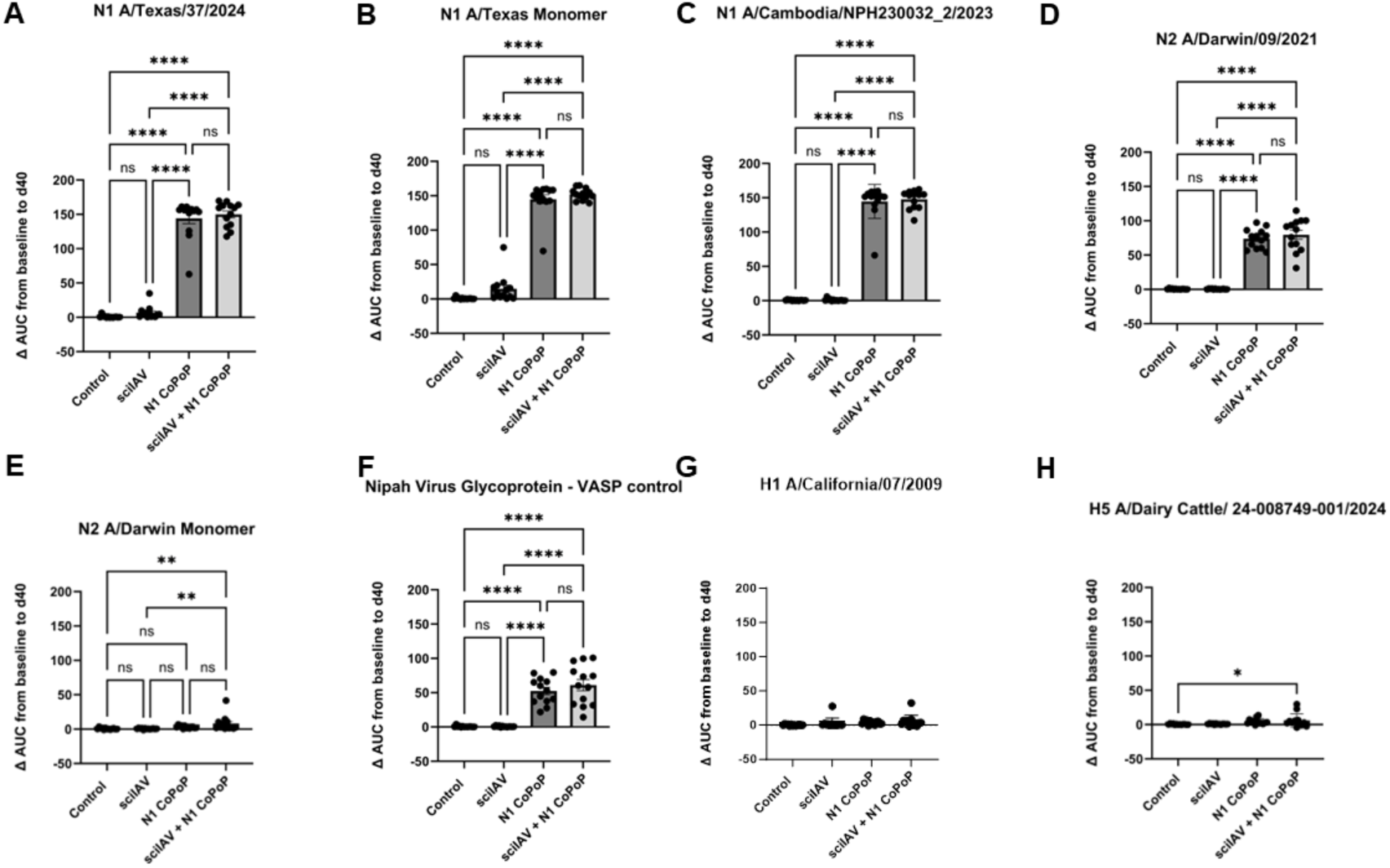
Difference in binding of serum IgG antibodies. Serum was harvested from blood collected from mice on day 0 and day 40 relative to vaccination with pH1N1 sciIAV on day 0 and/or two (day 14 and day 28) inoculations of CoPoP decorated with H5N1 NA proteins (1.5 μg NA A/Cambodia/NPH230032_2/2023 H5N1 plus 1.5 μg NA A/Texas/37/2024 H5N1). Serum from mice (N=13/treatment) was diluted 1:200 and 1:2,000 and assayed by ELISA in duplicate against each antigen shown. For determining the change in binding, results were calculated as area under the curve (AUC) and the difference between day 40 and day 0 was used in ordinary one-way ANOVA with Tukey’s multiple comparison test.

All mice that were mock-treated and infected with A/Texas/37/2024-Nluc, lost 25% of their initial bodyweight by 7 to 10 days after challenge (**Figure 2A**) and were humanely euthanized. Mice that were immunized with either pH1N1 sciIAV or N1-CoPoP, alone or in combination, were protected from severe weight loss and had 100% survival (**Figure 2A and B, respectively**). Mice that were administered N1-CoPoP either alone or in combination with pH1N1 sciIAV had higher body weights than mice given a single dose of pH1N1 sciIAV, although this difference did not reach significance (*p* > 0.05).

**Figure 2.**
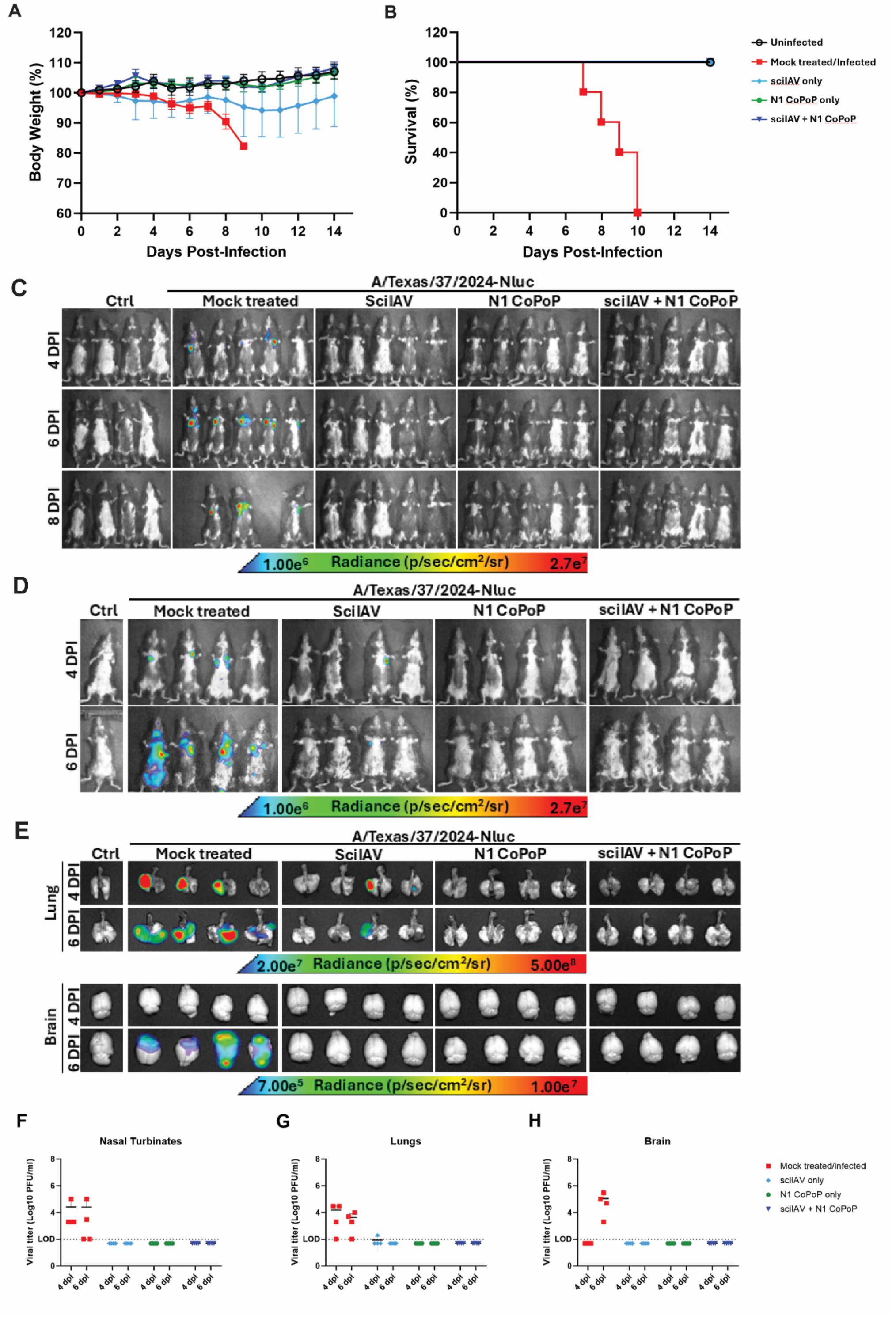
Protective efficacy of SciIAV, N1 CoPoP and SciIAV+N1 CoPoP in C57BL/6 Mice following infection with HPAI A/Texas/37/2024-Nluc virus. Female mice (4 mock infected and 5/treatment group) were vaccination with pH1N1 sciIAV on day 0 and/or two (day 14 and day 28) inoculations of CoPoP decorated with N1 proteins (1.5 μg NA A/Cambodia/NPH230032_2/2023 H5N1 plus 1.5 μg NA A/Texas/37/2024 H5N1) and then challenged with 1,000 PFU of A/Texas/37/2024-Nluc H5N1. Mice were followed for up to 14 days post infection for Body weight (A), survival (B). Body weight is expressed as a percentage of the initial body weight measured on the day of challenge (day 0). All mice that lost more than 25% of their body weight were euthanized. NanoLuc (Nluc) expression was monitored in vivo for body weight groups at 4, 6, and 8 DPI (C) and necropsy groups at 4 and 6 DPI (D), and ex vivo in the lungs and brains at 4 and 6 DPI (E). Ctrl = control. Radiance, expressed as photons per second per square centimeter per steradian (photons/s/cm²/sr), is indicated on the corresponding heatmaps. Four mice per treatment were humanely euthanized at 4 and 6 dpi and (F) nasal turbinates, (G) lungs, and (H) brains were collected and homogenized to evaluate viral titers. The limit of detection (LOD) was 100 PFU/ml. Tissues with undetectable viral titers were given a value of 50 for graphical and statistical representations.

An in vivo imaging system (IVIS) was used to visualize the distribution of bioluminescent signal in the tissues and whole mice challenged with A/Texas/37/2024-Nluc H5N1. In the mice groups used for mortality (n = 4 for mock treated; n = 5 for all treated mice), bioluminescence was measured on days 4, 6 and 8 following viral challenge (**Figure 2C**). Only mice that were mock-treated and virally challenged had detectable bioluminescence. Three mice in this group had evident signal at 4 dpi but all five had substantial signal by 6 dpi which was maintained in the surviving three mice at 8 dpi. No bioluminescence was detected in mice inoculated with a single dose of pH1N1 sciIAV or two doses of N1-CoPoP either alone or in combination with pH1N1 sciIAV (**Figure 2C**). In the mice groups used for necropsy, animals that were mock-treated and infected showed the most intense bioluminescence (**Figure 2D**). The amount of reporter

signal was high in the lungs of three mock-treated and infected mice by 4 dpi (**Figure 2E, upper panels)**. By 6 dpi, the signal had spread beyond the lungs in all four mice and was also evident in the brain (**Figure 2E, lower panels**). In the group of mice administered with pH1N1 sciIAV and then challenged with the reporter virus, two mice showed areas of bioluminescence in the lungs by 4 dpi **Figure 2E**). By 6 dpi, signal had diminished and was only detectable in the lungs of one mouse administered pH1N1 sciIAV alone. Mice inoculated with two doses of N1-CoPoP either alone or in combination with pH1N1 sciIAV showed no bioluminescence at any time point. These results suggest that both pH1N1 sciIAV and N1-CoPoP protect against severe infection with HPAI H5N1.

Viral titers were measured in nasal turbinates, lungs and brains of infected mice (n = 4 per treatment) on 4 and 6 dpi. Only mice that were mock-treated and infected had detectable virus in the nasal turbinates, 4 and 6 dpi (**Figure 2F**). Similarly, viral titers in lungs were significantly higher in mock-treated and infected mice 4 dpi than mice in other treatment groups (*p* = 0.008; **Figure 2G**). Only one mouse inoculated with pH1N1 sciIAV had detectable virus in lungs at 4 dpi. By 6 dpi, viral titers in lungs of mock-treated and infected mice had declined (*p* = 0.005) but was not different from lung viral titers of mice from other treatments (*p* > 0.05). No mice had detectable virus in the brain at 4 dpi (**Figure 2H**). However, mice that were mock-treated and infected had detectable viral titers in the brain at 6 dpi. All mice in other treatment groups had undetectable viral titers in the brain. This provides further evidence that either pH1N1 sciIAV or CoPoP decorated with N1 proteins can provide protection against HPAI H5N1 infection.

## Discussion

Although the current pandemic risk for HPAI H5N1 is rated as low, with continued circulation in mammalian reservoirs, that could change in the future. With the right mutations or an ominous rearrangement event that has similarly preceded every influenza pandemic in human history, producing an effective vaccine, is rapidly needed to reduce the potential impact to human life. The use of sciIAV has the advantage of stimulating immune memory against targets that are highly conserved between influenza viral strains. Internal viral proteins like nucleoprotein (NP), polymerase basic 1 (PB1) protein, and non-structural 1 (NS1) protein following MHC-I peptide presentation on infected cells can lead to T cell engagement and eventually establishment of tissue resident memory which can be rapidly mobilized following another influenza infection to clear infected cells. In our study, mice that received pH1N1 sciIAV were protected against H5N1 infection despite the vaccine strain being based on pH1N1. This heterosubtypic protection is typical for animals inoculated using sciIAV and subsequent viral challenge ^20,21^. Although we did not specifically measure T cell activity in this study, it has been investigated by others ^9,10^. Baker et al. ^9^ saw increases in CD8+ NP-tetramer+ cells in mice following prime and boost with 10^5^ PFU sciIAV which was similar to the increase in mice, 10 days following infection with wild-type virus. Mice given either S-FLU or CLEARFLU sciIAV had similar increases in CD8+ T cells in lungs and spleen responding to NP as compared to mice given control media ^10^. Because sciIAV is limited to a single round of replication in non-complementing systems, there is only finite engagement of the humoral immune system leading to little or no antibody generation against the dominant surface HA or NA glycoproteins (**Figure 1**). Because we only measured IgG in serum, this might not be unexpected given the intranasal route of inoculation, mucosal IgA may be more likely produced ^22^. Furthermore, we did not test for antibodies targeting other viral proteins like NP which may have contributed to viral clearance. It is not known if preexisting immunity to influenza either through previous infection or vaccination may influence T cell responses following vaccination with sciIAV. Compared to infection, sciIAV can evoke CD8+ T cell memory with more diverse TCR repertoire ^20^. Having greater diversity is beneficial for reacting to greater viral variation and limits successful escape mutations. Previous immunity may down select this diversity, resulting in reduced breadth of protection compared to the heterosubtypic protection we show in this study.

Recombinant proteins of the pandemic strain’s HA and NA can be utilized in a more targeted vaccine. Generation time can be measured in weeks rather than the months required for traditional egg based live attenuated or inactivated virus vaccines. Moreover, NA is more highly conserved than HA. For instance, the NA of pH1N1 has ∼90% homology to the NA of A/Texas/37/2024 H5N1, the strain used in this experiment for challenging mice. Because of the higher conservation of NA, a vaccine based on this protein could be utilized to provide broader protection than the HA focused seasonal influenza vaccines ^23^. The NA of influenza is a sialidase enzyme which primarily functions to release viruses that have engaged with sialic acid via hemagglutinin either from mucins displaying decoy sialic acid or more importantly from the cell surface so viral progeny can be disseminated to infect other cells. To be active, NA needs to be in the proper configuration as a tetramer^24^. Although both monomers and tetramers are immunogenic, only the tetramer seems to induce protective antibodies against viral challenge ^25,26^. Antibodies that inhibit the enzymatic activity of NA protein by either directly interacting with the active site or sterically hindering the active site would be best for neutralization and NA based vaccines that can induce these antibodies would certainly be ideal. Antibodies that target the NA active site also would provide the broadest protection because this site is highly conserved between influenza A and B viruses ^27^. The more epitopes an antibody binds to outside of the active site, however, due to their variability would narrow the protective breadth to viruses within an NA subtype, clade or even viral strain. Optimizing NA active site targeting antigens for vaccines remains a challenge.

Utilizing lipid nanoparticles like the CoPoP in this study improves the immunogenicity of the recombinant proteins and more closely mimics presentation of these proteins on the surface of viral particles. Similarly, self-assembling protein nanoparticles have been used for recombinant protein to present these antigens as an ordered, multimer including ferritin (24-mer ^28^), E2p or I3-01v9 (both 60-mer ^29^) using Spy tag/Spy Catcher ^30^ or Fagy tag ^28^ to allow easy plug-and-play conjugation to multimers. Decorating the outside of CoPoP liposomes with recombinant protein is even simpler, requiring only mixing of CoPoP liposomes and proteins containing a His-tag, already required for protein purification.

CoPoP liposomes decorated with recombinant influenza glycoproteins has been evaluated as a potential vaccine alternative ^17,31,32^. When only the NA of A/Wisconsin/588/2019 H1N1, A/Darwin/6/2021 H3N2, and B/Austria/1359417/2021 Victoria were decorated on the CoPoP liposome and used to inoculate mice, animals challenged with pH1N1 or A/Hong Kong/1/1968 H3N2 were not completely protected and some or all mice lost more than 25% of their body weight ^17^. This was attributed to the lack of homology between inoculation and challenge strains. Mice inoculated with NA of H1N1 and the NA of influenza B also had reduced viral loads in lungs 4 days after challenge. In this current study, inoculating with N1-CoPoP liposomes resulted in undetectable viral titers on days 4 and 6 after challenge with A/Texas/37/2024-NLuc H5N1. All mice that were inoculated with N1-CoPoP essentially maintained or gained weight after viral challenge and survived until the end of the experiment. This is likely the result of challenge virus matching to one of the NA proteins decorating the CoPoP liposome. In another study where clade 2.3.4.4b A/Jiangsu/NJ210/2023 H5N1 HA and NA recombinant proteins were used to decorate CoPoP liposomes, mice that received the N1-CoPoP liposomes were not completely protected (7 of 10 survived) after challenge with RTH/2022 2.3.4.4b H5N1 whereas those mice inoculated with CoPoP liposomes containing either the HA or both the HA and NA proteins were fully protected ^32^. This suggests that for NA, NA-CoPoP provide better protection when matched to the challenge virus.

Vaccinating with either sciIAV or N1 decorated CoPoP protected mice after HPAI H5N1 challenge. Using sciIAV seems to provide protection even from heterosubtypic challenge and occurred without significant humoral immune system engagement. Inoculation with N1-CoPoP, however, does induce N1 antibodies which may be the main route of protection against HPAI H5N1 challenge in N1-CoPoP immunized mice. Both sciIAV and CoPoP liposomes, alone or in combination, deserve further consideration as potential rapid implementation vaccine alternatives before the next influenza pandemic.

## Materials and Methods

### Biosafety

Experiments involving HPAI H5N1 were conducted in biosafety level 3 (BSL3) and animal BSL3 (ABSL3) high containment facilities at Texas Biomedical Research Institute. All procedures were conducted in accordance with protocols approved by the Texas Biomedical Research Institutional Committee (IBC #21-017) and Animal Care and Use (IACUC # 1785 MU) committees.

### Cells

Parental Madin-Darby canine kidney (MDCK) and human embryonic kidney (HEK293T) cells were used to generate and propagate stocks of recombinant viruses. Cells were cultured in Dulbecco’s modified Eagle medium (DMEM) (Invitrogen, USA) supplemented with 10% fetal bovine serum (FBS) and 1% PSG (penicillin, 100 U/mL; streptomycin, 100 μg/mL; L-glutamine, 2 mM) at 37°C with 5% CO_2_. MDCK cells constitutively expressing pH1N1/E3-HA were previously described ^9^.

### Single-cycle infectious influenza pH1N1

The single-cycle infectious pandemic influenza A/California/04/09 H1N1 (sciIAV pH1N1) was previously described ^9^. Briefly, seven pH1N1 ambisense plasmids (pDZ PB2, PB1, PA, NP, NA, M, and NS) ^33^, pPolI HA(45)GFP(80) ^34^, and pCAGGS pH1N1/E3-HA were transfected into 1:1 co-cultures of 293T and MDCK cells. Sixteen hours post-transfection, the media were replaced with DMEM containing 0.3% BSA, 1% PSG, and 1 μg/ml of TPCK-treated trypsin (Sigma). Virus was scaled up using cleared cell culture supernatant into MDCK cells expressing pH1N1/E3-HA and then titrated by plaque assay on MDCK cells expressing pH1N1/E3-HA.

### CoPoP assembly

The NA for A/Cambodia/NPH230032_2/2023 H5N1 (EPI2419979, amino acids 63-449) and A/Texas/37/2024 H5N1 (EPI3171486 amino acids 76-449) were synthesized by GenScript (Piscataway, NJ) with a 7x His-Tag, Avi-Tag and a tetramirization domain either from tetrabrachion (TB) from *Staphylothermus marinus* (A/Cambodia) or from human vasodilator-stimulated phosphoprotein (VASP; A/Texas) fused at the N-terminal. Lipid-porphyrin nanoparticles were synthesized as previously described ^35–37^. CoPoP liposomes were prepared as previously described ^38^. In short, the lipid bilayer included 1,2-dioleoyl-sn-glycero-3-phosphocholine (DOPC) and cholesterol for stabilization along with synthetic MPLA (PHAD-3D6A) a TLR4 agonist and QS-21 an inflammasome activator. The PHAD-3D6A and QS-21 each made up approximately 22.2% of the total liposome by weight prior to the addition of neuraminidase. To decorate the liposome, a 4:1 ratio of CoPoP liposome to total neuraminidase (50:50 A/Cambodia/NPH230032_2/2023 and A/Texas/37/2024) was co-incubated at 37°C for three hours and then stored overnight at 4°C. Incorporation of the NA proteins was confirmed by using 2 μg, heated to 95°C for 10 minutes in reducing buffer and resolving on a 4-12% Bis-Tris acrylamide gel by electrophoresis. The gel was stained with ProtoBlue Safe (National Diagnostics; Atlanta, GA) so that soluble and particle incorporated NA could easily be determined.

### *in vivo* studies

Female C57BL/6 mice (5 weeks of age, 13 mice per group, except mock-treated/mock-infected which included only 5 mice) were purchased from the Jackson Laboratory and maintained in the animal facility at Texas Biomedical Research Institute under specific, pathogen-free conditions and ABSL3 containment. After an acclimation period, mice were anesthetized using gaseous sedation in an isoflurane chamber for retro-orbital blood collection and intranasal (IN) inoculation with sciIAV (10^6^ PFU/mouse) in the two groups of mice receiving this treatment (sciIAV and sciIAV + CoPoP). Serum was harvested from collected blood and stored at −20°C. At 14 and 28 days later, mice in the CoPoP and sciIAV + CoPoP groups received intramuscular (IM) injections with 50 μl of CoPoP-NA (1.5 μg NA A/Cambodia/NPH230032_2/2023 H5N1 plus 1.5 μg NA A/Texas/37/2024 H5N1). On day 40, mice were anesthetized, blood collected for harvesting serum, and all but the five mock-treated/mock-infected mice were challenged IN with 1,000 PFU/mouse of the previously described influenza A/Texas/37/2024 H5N1 expressing Nluc ^39^. Five (or four, mock-uninfected) mice per group were followed for 14 days after viral challenge for changes in body weight and survival. Animals in these groups were also used *for in vivo* imaging on 4, 6, and 8 dpi (described below). The remaining mice in each group (8 mice per treatment group and 2 uninfected) were utilized for determining viral replication in nasal turbinates, lungs, and brain. Prior to euthanasia on either 4 or 6 dpi, mice were subjected to *in vivo* imaging as described below. Four mice from each experimental group and one of the uninfected mice were euthanized by administration of a lethal dose of pentobarbital sodium (Fatal Plus, Vortech Pharmaceuticals) followed by exsanguination on 4 and 6 dpi. Tissues were surgically extracted and homogenized in 1 mL of PBS using Precellys tissue homogenizer (Bertin Instruments, Rockville, MD, USA). Virus titers (PFU/mL) were determined by standard plaque assay in MDCK cells ^40^. Data representation of mean titers and determination of significant differences between viral titers of treatment groups were determined by One-way ANOVA and Tukey’s multiple comparisons test using GraphPad Prism (v9.4.1.458).

For *in vivo* imaging, mice were anesthetized on imaging days and then injected retro-orbitally with 100 μL of the Nano-Glo luciferase substrate (Promega, US) diluted to 1:10 in PBS. *In vivo* Nluc imaging was immediately performed using the IVIS Spectrum multispectral imaging system (Revvity, Inc.) ^39^. Bioluminescence data were acquired and analyzed using the Aura program (AMI Spectrum).

### Enzyme-linked immunosorbent assay (ELISA)

ELISA plates (Nunc MaxiSorp; Thermo Fisher Scientific, Rochester, NY, USA) were coated with recombinant NA, HA, or Nipah Virus glycoprotein (VASP tetramerization control) at 2 µg/mL in DPBS with calcium and magnesium (Corning Mediatech) ^41^. All recombinant proteins used were synthesized at GenScript in TurboCHO™ cells except the HA for A/California/07/2009 H1N1 and A/Texas/37/2024 H5N1 which were provided by BEI Resources. Serum was heat inactivated at 65°C for 30 minutes and diluted in DPBS at 1:200 and 1:2,000, and binding was detected with HRP-conjugated anti-mouse IgG (Jackson ImmunoResearch, West Grove, PA, USA). After washing with PBS-Tween (0.05%), SureBlue TMB Microwell Substrate (Sera Care Lifesciences, Inc., Milford, MA) was added. After 10 minutes incubation, color development was stopped with the addition of 1N HCl and optical density at 450nm was read on a Versa Max plate reader (Molecular Devices, LLC, San Jose, CA). For area under the curve (AUC) calculations, the inverse of the dilution was multiplied by 10,000 for the x value and the OD at 450 nm was used as the Y. AUC was determined in Graphpad Prism (Ver. 10). The difference between day 40 and day 0 AUC was used in ordinary one-way ANOVA with Tukey’s multiple comparison test. Treatment differences were considered significant with a *p* ≤ 0.05.

## Acknowledgements

This work was partially supported by a grant from the American Lung Association (ALA) to L.M-S, a Texas Biomedical Research Institute Forum Award to A.M.E., and a Douglass foundation award to R.S.B..

The following reagents were obtained through BEI Resources, NIAID, NIH: [HA for A/California/07/2009 and A/Dairy Cattle/ 24-008749-001-original/2024], catalog numbers NR-51668 and NR 59816, respectively. Microsoft Copilot was used to help summarize literature used in the introduction and was checked for accuracy by M.P.

## Conceptualization

J.J.K., M.R.W., J.F.L and L.M.-S.; Methodology: M.S.P, R.S.B. Q.L., R.A.E., A.M.E and L.M.-S.; Data collection and interpretation: M.S.P, R.S.B., R.A.E., J.J.K., and L.M.-S.; Funding acquisition and resources: J.J.K., A.M.E. and L.M.-S.; Writing—original draft preparation: M.S.P and R.S.B.; Writing—review and editing: all authors have read and agreed to the published version of the manuscript.

